# Rice-Fish Co-Culturing for Sustainability, Food Security, and Disease and Poverty Reduction

**DOI:** 10.1101/2025.11.14.688524

**Authors:** Emily K. Selland, Nicolas Jouanard, Amadou Guisse, Momy Seck, Andrea J. Lund, David López-Carr, Alexandra Sack, Louis Dossou Magblenou, Giulio De Leo, Molly J. Doruska, Christopher B. Barrett, Jason R. Rohr

## Abstract

Over 140 million households cultivate rice worldwide. However, rice production contributes to exceedances of planetary boundaries, such as freshwater use and biogeochemical flows. Additionally, rice-farming families may face increased risk of acquiring schistosomiasis, a parasitic disease transmitted by freshwater snails that contributes to reinforcing cycles of poverty and disease. Using data from 405 households in rural Senegal, we show that children in rice-farming households had higher *Schistosoma mansoni* prevalence and *S. haematobium* intensities than non-farming peers. To address this environmental-health challenge, we integrated native Nile Tilapia (*Oreochromis niloticus*) and African Bonytongue (*Heterotis niloticus*) into rice fields. The fish thrived, suppressed insects and snail pests, improved soil nutrients, and boosted rice yields by >25% with a net benefit of 1,805–3,415 USD/ha/year (benefit-to-cost ratio = 7.42). Hence, low-input rice–fish co-culturing offers a scalable planetary health solution that simultaneously advances sustainability, nutrition, health, and rural livelihoods.

**Résumé:** Plus de 140 millions de foyers à travers le monde cultivent du riz. Cependant, cette production exerce une pression importante sur les ressources naturelles, notamment les réserves d’eau douce. Par ailleurs, les ménages pratiquant la riziculture peuvent être plus exposés au risque de bilharziose, une maladie parasitaire transmise par des escargots d’eau douce, qui contribue au cycle de pauvreté. À partir de données collectées auprès de 405 ménages dans des villages sénégalais, nous montrons que les enfants issus de familles pratiquant la riziculture présentent une prévalence plus élevée d’infection à *Shistosoma mansoni* et des intensités d’infection à *S. haematobium* plus importantes que ceux issues de familles ne cultivant pas le riz. Pour remédier à cela, nous avons intégré deux espèces locales de poissons, le tilapia du Nil (*Oreochromis niloticus*) et l’Hétérotis (*Heterotis niloticus*), dans les rizières, afin de réduire les population d’escargots hôtes, améliorer les rendements agricoles et augmenter les revenus. Ces poissons ont montré une bonne croissance, ont contribué à la régulation des insectes et des escargots nuisibles, ont enrichi le sol en nutriments, et accru les rendements rizicoles de plus de 25%, avec un bénéfice net estimé entre 1 805 et 3 415 USD/ha/an (ratio bénéfice-coût = 7,42). Ainsi, la co-culture riz-poisson à faibles intrants constitue une solution reproductible en faveur de la santé publique, permettant de concilier durabilité, nutrition et amélioration des moyens de subsistance en milieu rural.

## Main

The interdependence between human health and the health of the environment has become increasingly recognized. “Planetary Health” and “One Health” frameworks are being adopted to understand the ways in which reversing or preventing environmental degradation can simultaneously benefit animal and human health and environmental sustainability^1^. Efforts to control and understand both non-communicable and communicable diseases have revealed an increasingly evident connection between environment and human health^2–8^. One example is rice-fish co-culturing, which has been proposed for reducing widespread under- and malnutrition and increasing the sustainability of rice production.

Rice is a staple food and the primary source of daily calories for much of the world’s population^9,10^, and its demand is increasing, especially in Africa^11^— the continent with the fastest growing population^12^. However, rice farming contributes to the crossing of several planetary boundaries, such as land-system change, freshwater use, and nitrogen flows^13,14^. Growing rice and fish together might help mitigate these impacts by improving efficient land and water use, while also providing economic stability^15–17^ and nutritional diversification for rice farming households^18,19^, who report higher income levels, fish consumption^20,21^, and rice yields^16^. Additionally, co-culturing often leads to reduced fertilizer and pesticide inputs as fish can increase nutrient availability to crops^22–25^ and consume or compete with pests of rice plants^26^. Despite being common practice in Asia, there have been only a few attempts to test rice-fish co-culturing in Africa^27–30^.

Here we argue that rice-fish co-culturing may have the additional potential benefit of reducing schistosomiasis, a chronic disease that afflicts over 250 million people, >90% in Africa^31,32^. Schistosomiasis is caused by parasitic *Schistosoma* spp. flatworms released from freshwater snails that penetrate human skin and contributes to trapping rural communities in poverty^31,33,34,35^. As rice is typically grown in water, rice paddies can harbor snails, thus exposing rice farmers to *Schistosoma* parasites^36^. However, with a few exceptions^37,38^, there is a paucity of rigorous studies that quantify the occupational risk of acquiring schistosomiasis for rice farmers in Africa, although recent work detected snails shedding schistosomes in rice fields^39^. The idea of using natural enemies of pathogens and their vectors to curb the transmission of environmentally mediated disease is gaining increasing attention, and an occupational risk of schistosomiasis could be mitigated by adding fish to rice fields that either consume or compete with snails, as biological control has been effective at reducing *Schistosoma* transmission in other endemic regions^40–43^. The use of carp to control non-disease-vector snails is common in Southeast Asian rice fields^44,45^, but carp are not native to Africa and their introduction could damage surrounding ecosystems^46^. Here, we investigate co-culturing of native fish and rice in West Africa.

Developing snail biocontrol in African rice fields is important for several reasons^43^. First, although the drug praziquantel can cure existing infections, it does not prevent reinfection. Therefore, treated people may acquire new infections when they return to snail-infested waters, threatening disease elimination goals. Thus, interventions that target snails have been increasingly called upon to better control schistosomiasis^47–50^. Second, common snail control approaches like molluscicides have temporary effects as snails can recolonize habitats once the chemical degrades. Additionally, molluscicides are toxic to non-target species and make water unsafe for human consumption^51^.

In the lower Senegal River valley (SRV), the Diama Dam benefits agricultural producers through desalination and irrigation, leading to substantial increases in irrigated rice farming^19,52^. Associated increases in irrigation and fertilizer runoff also augmented snail habitat and food (algae), causing an eruption of schistosomiasis^2,52^, among other socioeconomic issues^19^. The dam also reduced snail predators, such as prawns and fish, releasing snail populations from natural biological controls^53^ and further exacerbating schistosomiasis. Thus, as rice production proliferates in Africa^11^, there is an increasing potential occupational risk of schistosomiasis unless effective snail control is implemented in rice fields.

Here, we first test the hypothesis that rice farmers face an occupational schistosomiasis hazard by assessing whether children of rice farming households in the SRV have higher rates of *Schistosoma* infections than non-rice farming households. Second, we test the hypothesis that adding native fish predators (*Heterotis niloticus* – African Bonytongue, which predate snails^39^) and competitors (*Oreochromis niloticus* – Nile Tilapia, which consume algae^23^) of snails to rice fields would reduce snails and improve sustainability of rice farming by increasing rice and fish production with a low external input sustainable agriculture (LEISA) system (i.e., fish feed not added to fields)^39^. To identify potential mechanisms for any changes in rice production, we quantify insect abundance and soil and water nutrient levels. Finally, we assess whether rice-fish co-culturing is profitable in the SRV through an economic benefit-cost analysis. By testing a LEISA, ecologically grounded system using native species in Senegal’s schistosomiasis-endemic rice fields, we aim to evaluate whether rice-fish co-culturing can simultaneously suppress disease vectors, enhance agricultural productivity, and improve the livelihoods of farmers. This work sets the stage for an integrated, scalable intervention that aligns food production with environmental health and infectious disease control in similarly burdened regions.

### Is there an Occupational Hazard of Rice Farming?

To test whether rice farming is an occupational hazard for schistosomiasis, we assessed whether the number of hectares of rice farmed by a family is a predictor of schistosomiasis transmission risk using a household survey and parasitological data across schoolchildren in the SRV. The parasitological data were gathered from villages where schistosomiasis treatment with praziquantel was not known to be performed the prior year for the majority of villages, such that a child’s worm burden largely reflects an accumulation of rice-farming effects over 1 or more years^54^. Many village-, household-, and individual-level variables have been shown to affect schistosomiasis levels in this region, such as the location of a village on the river or lake^2,55^, amount of village irrigated agriculture (total reported hectares of cultivated land fed through irrigation)^56^, availability of safe water^57^, household occupation (as children may assist with certain jobs)^58^, and age and sex of the tested child^59,60^. Therefore, to isolate the effect of household rice farming on *S. mansoni* and *S. haematobium* prevalence and intensity, we controlled for these hierarchical effects in our statistical models (see Methods).

Prevalence of *S. mansoni* was generally lower than that of *S. haematobium* within our sample (Figure 1A). For *S. mansoni* (cause of intestinal schistosomiasis), prevalence was positively associated with the age of the child and village located on the lake relative to the river and was negatively associated with the amount of village irrigated agriculture and household use of safe water for laundry (Table 1). After controlling for these effects, we found a positive association between infection prevalence in children and the number of hectares of rice that their household farmed (Table 1; Fig 1A). However, the association of *S. mansoni* infection intensity (measured by average egg output) with rice cultivation was not significant (Table 1; Fig 1B). For *S. haematobium* (cause of urogenital schistosomiasis), infection intensity was associated with rice cultivation, signaling greater worm burdens in children in rice farming households (Table 1; Fig 1B). The relationship was not significant for infection prevalence (Table 1; Fig 1A), perhaps because prevalence was so high (on average 68.9%) that there was little room for rice farming to increase prevalence (i.e., a ceiling effect). Hence, both *S. mansoni* and *S. haematobium* had some positive association with the level of household rice cultivation, suggesting an occupational risk of both intestinal and urogenital schistosomiasis for children of rice farming families. This is consistent with previous work showing that rice farming and household irrigated agriculture were risk factors for *Schistosoma* infections^36,37,56^.

**Figure 1.**
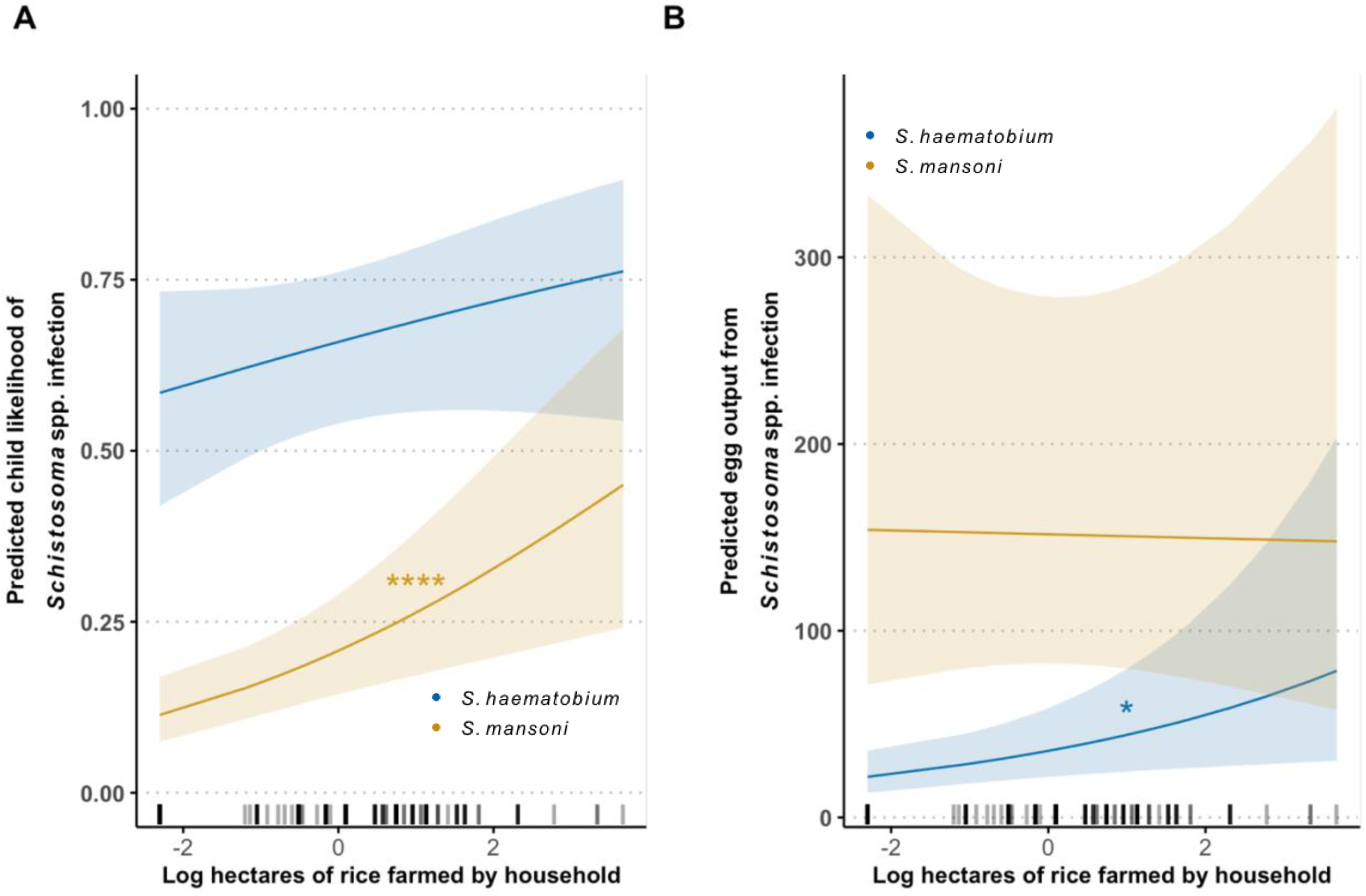
Household rice farming influences child infection outcomes. After controlling for individual-, household-, and village-level effects (see Table 1), there was a positive association between the hectares of rice that a household cultivated (as natural log+1 of household hectares) and (**A)** *Schistosoma mansoni* prevalence (X^2^ = 11.16, *p*<0.001), but not *S. haematobium* prevalence (X^2^ = 1.54, *p*=0.22) and (**B**) *S. haematobium* infection intensity (X^2^=5.86, *p*=0.015), but not *S. mansoni* infection intensity (X^2^=0.005, *p*=0.9) in the household’s school-aged children. Lines represent trends from predictions and shaded outlines represent 95% confidence intervals around the predictions. Rugs show each household replicate. Asterisks correspond to significance levels of *p*<0.1 (*) and *p*<0.001 (****).

**Table 1.** Regression model outputs for the best models (selected based on lowest AIC) for the prevalence and intensity of *S. mansoni* and *S. haematobium* among children. Household and village were included as nested random intercepts. All log values are natural log transformed.

| Dependent variable | Statistical test (Error distribution) | Predictors | Chi-square | Estimate (SE) | p value |
| --- | --- | --- | --- | --- | --- |
| Child *S. mansoni* infection | Regression (Binomial) | Child age | 10.75 | 0.2 (0.05) | 0.001 |
|  |  | Household hectares of rice (log) | 11.16 | 0.3 (0.09) | <0.001 |
|  |  | Village Irrigation Level | 35.34 | -0.02 (0.003) | <0.0001 |
|  |  | River v. Lake | 3.67 | 0.8 (0.4) | 0.06 |
|  |  | Piped Water | 6.98 | -0.6 (0.2) | 0.008 |
|  |  | Treated Village [Blocking] | 15.88 | 2.3 (0.6) | <0.0001 |
| Child *S. mansoni* infection intensity | Regression (Neg Binomial) | Household hectares of rice (log) | 0.005 | -0.007 (0.1) | 0.9 |
|  |  | Village Irrigation Level | 18.38 | 0.01 (0.009) | <0.0001 |
|  |  | River v. Lake | 3.16 | 5.7 (1.4) | 0.08 |
|  |  | Village Irrigation: River v. Lake | 13.72 | -0.03 (0.009) | <0.001 |
|  |  | Treated Village [Blocking] | 6.18 | -2.7 (1.1) | 0.01 |
| Child *S. haematobium* infection | Regression (Binomial) | Household hectares of rice (log) | 1.54 | 0.14 (0.1) | 0.22 |
|  |  | Village Irrigation level | 0.78 | 0.004 (0.006) | 0.38 |
|  |  | River v. Lake | 0.36 | 4.7 (2.0) | 0.55 |
|  |  | Village Irrigation: River v. Lake | 5.30 | -0.03 (0.01) | 0.021 |
|  |  | Treated Village [Blocking] | 2.43 | 1.4 (0.9) | 0.12 |
| Child *S. haematobium* infection intensity | Regression (Neg Binomial) | Household hectares of rice (log) | 5.86 | 0.22 (0.09) | 0.015 |
|  |  | Village Irrigation level | 0.013 | 0.0003 (0.003) | 0.91 |
|  |  | River v. Lake | 1.8 | 0.48 (0.4) | 0.18 |
|  |  | Treated Village [Blocking] | 2.98 | 0.85 (0.5) | 0.08 |

The occupational hazard of rice farming in the SRV could be further elucidated by addressing several gaps and limitations of the current work. First, measuring the infection likelihood in children is an indirect measurement of risk to rice farmers. We only had access to infection data from children, as they are the most commonly tested and vulnerable demographic^2,61–63^, but future studies should test this relationship among adults engaged in rice farming. Information on the water contact behavior of individuals at transmission sites (rice fields or other water bodies) and the inclusion of larger rice farms could also help elucidate the influence of rice on infection risk with respect to other transmission sites. Nonetheless, our work points towards rice farming in the SRV as an occupational hazard for schistosomiasis transmission worthy of control interventions.

### Can Native Fish Thrive in Rice Fields in this Arid Region?

Given an apparent schistosomiasis occupational hazard for rice farmers, we test whether rice-fish co-culturing might mitigate the risk of acquiring schistosomiasis while simultaneously increasing the productivity and environmental sustainability of rice. In 2023 and 2024, we stocked rice fields with fish during two cultivation seasons and maintained other fields as rice-monoculture control fields (i.e., without stocking fish). We recruited farmers, dug a trench (∼1 m deep) in each co-culture field to provide a deep-water refuge for fish, and added tilapia to all co-culture fields. Because of low African Bonytongue numbers, only a subset of fields receiving tilapia also received African Bonytongue during the first cultivation season.

Despite not being actively fed, fish thrived in rice fields without added inputs. During the first year, the fish were in treatment fields for an average of 81 days (SD =16.93), and tilapia were significantly larger at harvest than they were at stocking (*n*=9; Table 2; Fig 2C). African Bonytongue also grew in fields, with an average weight of 1,717 g at harvest compared to 454 g at stocking (*n*=6 fish; Table 2; Fig 2D). During the second trial, fish were in fields for only an average of 47.71 days (SD =14.33) and both tilapia and African Bonytongue did not meaningfully grow over this shorter season, likely due to low water conditions (see Methods). Further experiments should be conducted to further boost fish growth, such as by stocking mono-sex tilapia or adding fish predators to preclude competition from newly born fish, thus increasing the chances that stocked fish reach a commercial size^64^.

**Figure 2.**
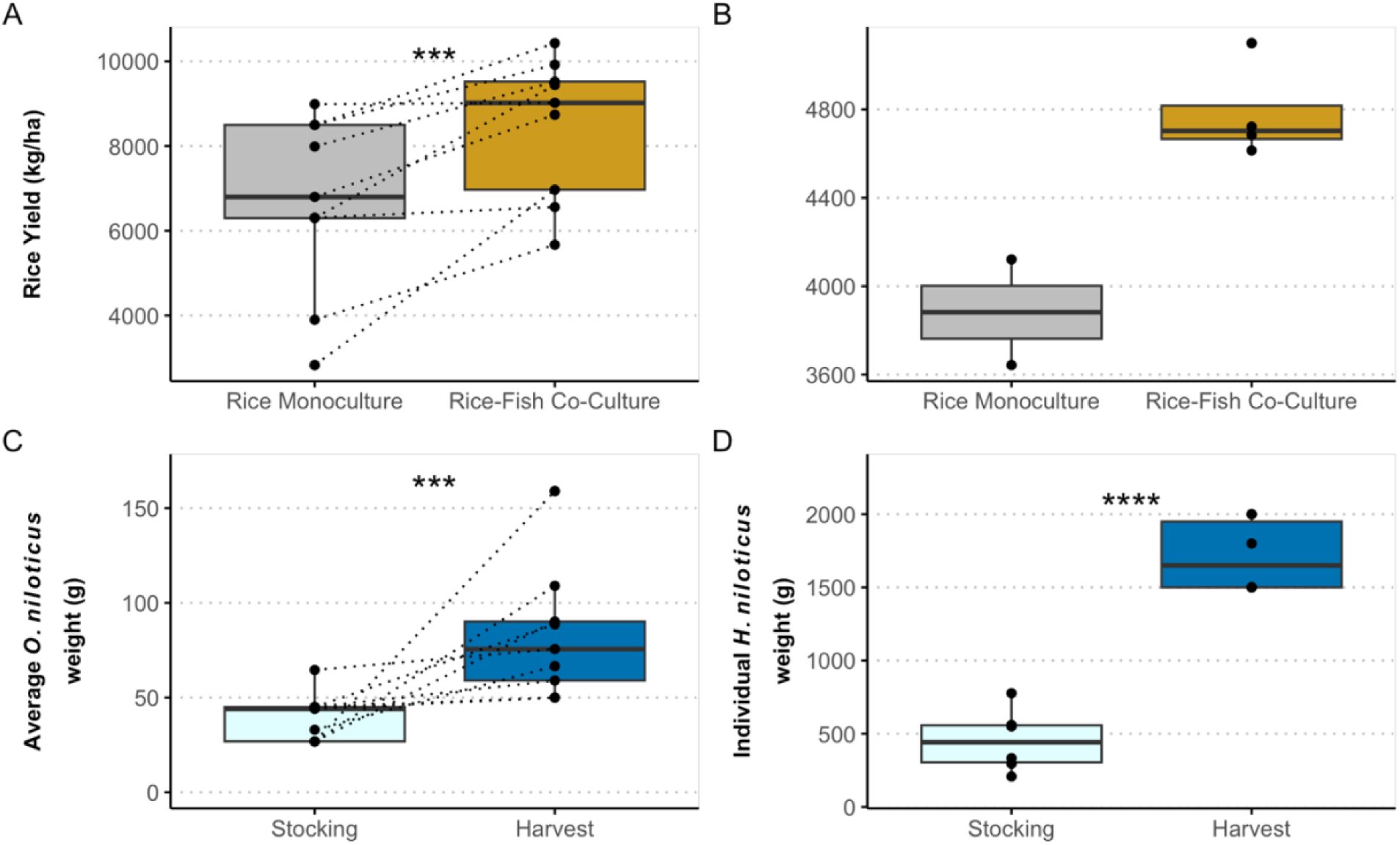
Rice-fish co-culture trials enhanced rice production and added fish production to rice fields. Rice yield (kg of rice per hectare) improved with rice-fish co-culturing during the **A.** first and **B.** second field experiments. **A.** During the first experiment, fields were paired to control for the management of individual farmers. Paired fields are represented with dashed lines. Every rice field that had fish (either *O. niloticus* or a polyculture of *O. niloticus* and *H. niloticus*) yielded rice equal to or greater than its paired control field. **B.** During the second year, fields were not paired, but the benefit of rice-fish co-culture on rice yield was still evident; however external water scarcity resulted in an underpowered test. Additionally, both **C.** *O. niloticus* (Nile Tilapia) and **D.** *H. niloticus* (African Bonytongue) were larger on average at harvest than at stocking, indicating growth in the rice fields in the absence of any fish feed. **C.** *O. niloticus* weight is calculated as the average weight per field, and the same fields at stocking and harvest are represented with dashed lines. **D.** *H. niloticus* weight is reported as the weight at stocking and harvest for each individual, but individuals cannot be matched. Box plots display average, interquartile range (IQR), and the 1.5xIQR. Asterisks correspond to significance levels of *p*<0.05 (**), *p*<0.01 (***), and *p*<0.001 (****).

**Table 2.** Results of statistical tests from rice-fish co-culture field experiments. Statistical test is listed for each endpoint. All log values are natural log transformed.

| Dependent variable | Statistical test<br>(Error distribution) | Predictors | Test statistic | Effect size or estimate (SE)<sup>a</sup> | *p* value |
| --- | --- | --- | --- | --- | --- |
| Rice yield (kg/ha) | *t*-test (2023) | Co-culture vs rice monoculture | *t* =4.21 | 1.40 | 0.0029 |
|  | *t*-test (2024) | Co-culture vs rice monoculture | *t* = 3.417 | 2.53 | 0.118 |
| *O. niloticus* growth | *t*-test | Stocking vs Harvest weight | *t* = 3.043 | 0.92 | 0.008<sup>b</sup> |
| *H. niloticus* growth | *t*-test | Stocking vs Harvest weight | *t* = 13.23 | 4.55 | <0.0001<sup>b</sup> |
| *Bulinus* abundance: | Regression (Neg Binomial)<sup>c</sup> |  |  |  |  |
| *truncatus/globosus* |  | *H. niloticus* presence | X<sup>2</sup> = 5.062 | -1.77 (0.8) | 0.012<sup>b</sup> |
|  |  | Number of *O. niloticus* harvested | X<sup>2</sup> = 4.11 | 0.72 (0.001) | 0.034 |
| *forskalii/senegalensis* |  | *H. niloticus* presence | X<sup>2</sup> = 0.17 | -0.40 (1.0) | 0.34<sup>b</sup> |
|  |  | Number of *O. niloticus* harvested | X<sup>2</sup> = 0.28 | -0.001 (0.002) | 0.60 |
| *Lymnaea* spp.<br>abundance | Regression (Neg Binomial)<sup>c</sup> | *H. niloticus* presence | X<sup>2</sup> = 1.88 | -1.25 (0.9) | 0.085<sup>b</sup> |
|  |  | Number of *O. niloticus* harvested | X<sup>2</sup> = 1.54 | 0.002 (0.001) | 0.22 |
| Insect Abundance | Regression (Neg Binomial)<sup>c</sup> | Co-culture vs rice monoculture | X<sup>2</sup> = 8.35 | -0.91 | 0.0037 |
|  |  | Depth | X<sup>2</sup> = 0.34 | -0.006 | 0.56 |
| Soil Samples | Regression (Gaussian)<sup>c</sup> |  |  |  |  |
| Carbon |  | Co-culture vs rice monoculture | X<sup>2</sup> = 7.49 | 0.39 (0.1) | 0.0062 |
| Total Nitrogen |  | Co-culture vs rice monoculture | X<sup>2</sup> = 8.88 | 0.032 (0.03) | 0.0029 |
| Available Phosphorus |  | Co-culture vs rice monoculture | X<sup>2</sup> = 1.66 | -10.23 (8.0) | 0.20 |
| Water Samples | Regression (Gaussian)<sup>c</sup> |  |  |  |  |
| Phosphorus (log) |  | Co-culture vs rice monoculture | X<sup>2</sup> = 2.97 | -0.35 (0.2) | 0.085 |
| Ammonia (log) |  | Co-culture vs rice monoculture | X<sup>2</sup> = 1.09 | 0.33 (0.3) | 0.30 |

<sup>a</sup> The estimate for *t*-tests is Cohen's *D*, whereas the estimate for any X<sup>2</sup> tests is the raw slope coefficient from the model.
<sup>b</sup> one-tailed test
<sup>c</sup> The test statistic for all regression analyses was a chi-square statistic.

### Can Native Fish in Rice Fields Reduce Snail Intermediate Hosts?

One mechanism by which the snail predator African Bonytongue could have grown in fields without supplemental feed is by consuming snails. To test this hypothesis, we compared snail abundance in fields stocked with African Bonytongue and tilapia relative to tilapia alone during our first trial. We measured the abundance of *Bulinus truncatus/globosus* (*B. t/g*), the primary snail hosts of *S. haematobium* in the region, *Bulinus forskalii/senegalensis* (*B. f/s*), minor contributors to *S. haematobium* transmission^65,66,2,55,67^ (but see^68^), and *Lymnaea* spp., the intermediate host for *Fasciola hepatica*, a liver fluke that causes considerable harm to livestock in the region^69^. Across both years, the rice fields had too few snails shedding *Schistosoma* cercariae and too few *Biomphalaria pfeifferi*, the intermediate host of *S. mansoni* in this region, to justify analyses.

Fields with African Bonytongue and tilapia had fewer total *B. t/g* snails – but not *B. f/s* - and tended to have fewer *Lymnaea* snails than those with only tilapia (Table 2; Fig 3A). This is consistent with laboratory trials showing that native African Bonytongue depredate snails^39^. During the second year, the low water level (see Methods) reduced overall snail presence/abundance in rice paddies (see Methods), and analyses of snail abundance were underpowered statistically. The lack of a reduction of *B. f/s* might be because these snails may be more associated with detritus than their *B. t/g* counterparts and therefore may be less likely to be consumed by African Bonytongue. The number of tilapia harvested from the field was positively associated with the abundance of *B. t/g*, likely because they both consume algal resources and thus fields with more nutrients likely supported more of both taxa (Table 2). This finding has implications for infected snails in rice fields, because greater algal resources could support greater transmission risk^70^. We encourage further work to assess the potential for fish-snail competition to reduce *Schistosoma* output from infected snails^71^.

**Figure 3.**
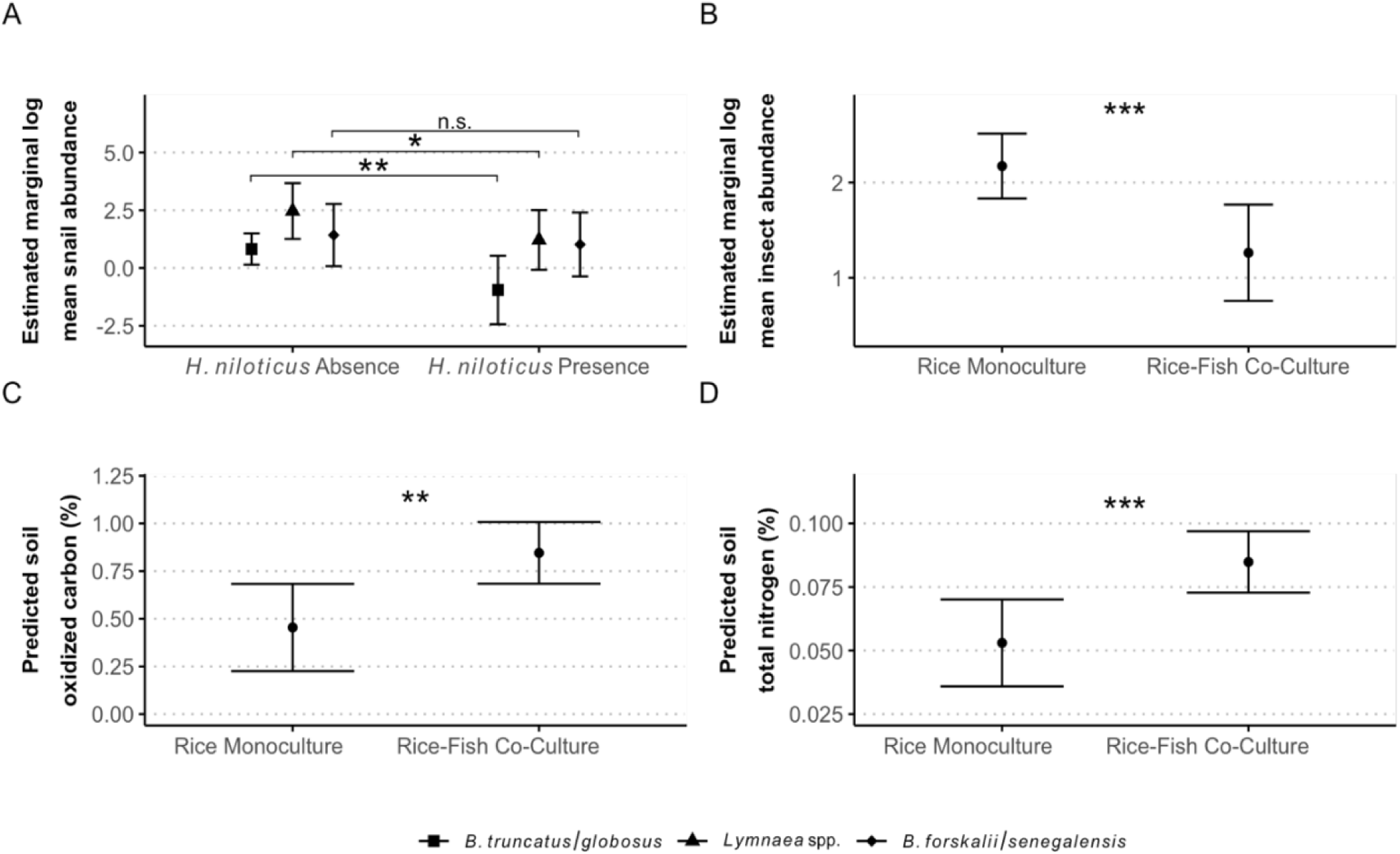
The presence of fish in rice fields altered snail and insect populations and soil nutrients. **A.** The presence of *Heterotis niloticus* in rice fields reduced the abundance of the main intermediate host of *Schistosoma haematobium*, *Bulinus truncatus/globosus* (*p*=0.0122; one-tailed), and *Lymnaea* spp. snails (*p*=0.085; one-tailed) that transmit *Fasciola hepatatica* to livestock in this region. **B.** Insect abundance (natural log transformed) was reduced in rice-fish co-culture fields compared to rice monoculture fields. Additionally, the predicted percentage of **C.** oxidized carbon and **D.** total nitrogen in soil were greater in fields where fish were present than absent. All figures show predicted or marginal means and associated 95% confidence intervals. Asterisks correspond to significance levels of *p*<0.1 (*), *p*<0.05 (**) and *p*<0.001 (***).

Despite not being detected during our study, *B. pfeifferi* have been previously found in rice fields in the SRV where intestinal schistosomiasis is endemic and its prevalence is associated with rice farming. This suggests that either temporal or spatial clustering may occur in the SRC, which has been shown to be important in previous work^72,73^. Future work should test the effects of rice-fish co-culturing on both *S. mansoni* and *S. haematobium* infection incidence in farmers in other areas of the SRV. Additionally, future studies should explore the use of other native fish species that might offer comparable or better biocontrol and profits^24,74–78^, such as catfish (*Clarias* spp.) that may both depredate and compete with host snails.

### Does Adding Fish to Rice Fields Affect Insects and Nutrient Availability?

We hypothesized that native fish might also reduce other invertebrates and influence nutrient levels in the ecosystem. To test this hypothesis, we measured insect abundance and soil and water nutrients in fields with or without fish additions during the second cultivation season (i.e., not a paired design, see Methods). Consistent with our hypothesis, rice fields with fish had fewer insects than those without fish (Table 2; Fig 3B), aligning with other work demonstrating that fish in rice fields can reduce insect pests^17,23,79^.

Next, we tested whether fish affected nutrient availability to rice by quantifying nutrients in the soil (carbon, total nitrogen, and available phosphorus) and water column (phosphorus, ammonia, and nitrate). Soil oxidized carbon, a measure of organic matter content, and total nitrogen were more abundant (Fig 3C, D), and available phosphorus was not different, in soil samples from co-cultured than monoculture fields (Table 2). The reduction in phosphorus in the water column (Table 2) was not significant, did not adversely affect rice yields (see below), and is contrary to the findings of other studies^80,16^. Nitrate levels were below detection limits (0.2 mg/L) for all water samples, and ammonia levels did not differ between treatments (Table 2), which is contrary to other work demonstrating fish presence can reduce ammonia^17,79,81^. The soil carbon improvements in co-culture fields indicate improved fertilization contributing to greater production^82^. Nitrogen, which is the most important nutrient associated with rice yield^83^, was low in the system.

Given that the addition of fish reduced insects and increased nutrient availability, it is likely that fish introductions might also reduce the need for insecticide and fertilizer use in fields. This has been demonstrated in Asian co-culturing systems^17,76^, but in our trials, farmers were asked to apply their normal fertilizer levels to both control and treatment fields. Future studies should test this hypothesis and quantify the effect of fish on the abundance of rice pests directly^84^. Notably, greenhouse gas emissions can vary by rice-animal system^76^, and we suggest that future work monitor rice field emissions with and without fish introductions.

### Does Adding Fish to Rice Fields Increase Rice Yields?

We expected that fish additions would increase rice yields^79,80^ and tested this in both paired and unpaired rice-fish co-culturing studies. As predicted, in our paired field study, rice yield was significantly greater in fields stocked with fish than those without fish (Table 2; Fig 2A), increasing average yields by 26.8%, from 6.68 to 8.47 tons/hectare (Cohen’s D = 1.40). In the second trial, rice yields were also greater in co-culture than monoculture fields that remained flooded throughout the growing season (*n*=6, Table 2; Fig 2B), although few control fields remained flooded throughout the season (*n*=2) leading to lower power in this year. Hence, in repeated trials, native fish additions increased rice yields. Greater rice yields in co-culture fields were expected^80^, however our test is the first in Senegal. We suspect that the greater yields were a product of greater nutrient availability and potentially fewer insect pests of rice^81^.

### Is Rice-Fish Co-Culturing Economically Viable in Senegal?

To test whether the observed fish growth in the absence of supplemental feed and the increased rice yields with co-culturing demonstrated a high economic return to rice-fish co-culturing, we conducted a cost-benefit analysis (see Methods). The primary cost of rice-fish co-culturing is a one-time investment in a fish refugia trench, and the main recurring cost is fingerling fish procurement. The primary benefits of co-culturing include improving rice yields by 1,793.5kg/ha compared to monoculture fields (See Fig 2A, B) and Nile tilapia production, with an average per-capita mass of 0.0806 kg at harvest (Supplemental Table 3). This results in an average of 59.6 kg of tilapia/ha in total and rice revenues improving by $479.04/ha/year. While tilapia’s average mass is lower than values typically reported in the literature under different environmental conditions^85^, we used mixed-sex tilapia which reduced average mass because of in-field reproduction and participating farmers were raising fish for the first time. Thus, these results suggest promise with room for improvement.

Based on the market value of Nile tilapia (Supplemental Table 3), the estimated range of fish and rice benefits is 2,345.52 - 3,774.64 USD/ha. Together, these revenues ensure that the initial trench investment of 1,000 USD/ha is recovered in the first year and the net benefit is 1,805.48 – 3,414.64 USD/ha/year, amounting to a median benefit-to-cost ratio of 7.42 and a range of ratios from 4.34 - 10.49. Hence, the average net benefit per capita for rice farming households is 379.16 - 717.07 USD/year. We additionally found that the distribution of rice yields in co-culture fields increased relative to monoculture fields (Fig 4), indicating that co-culture reduces the risk of very low rice yields, which will appeal to risk-averse farmers. While we suggest rice yield be monitored in further experiments, this provides a general result favoring rice-fish co-culture over monoculture than a simple comparison of mean rice yields.

**Figure 4.**
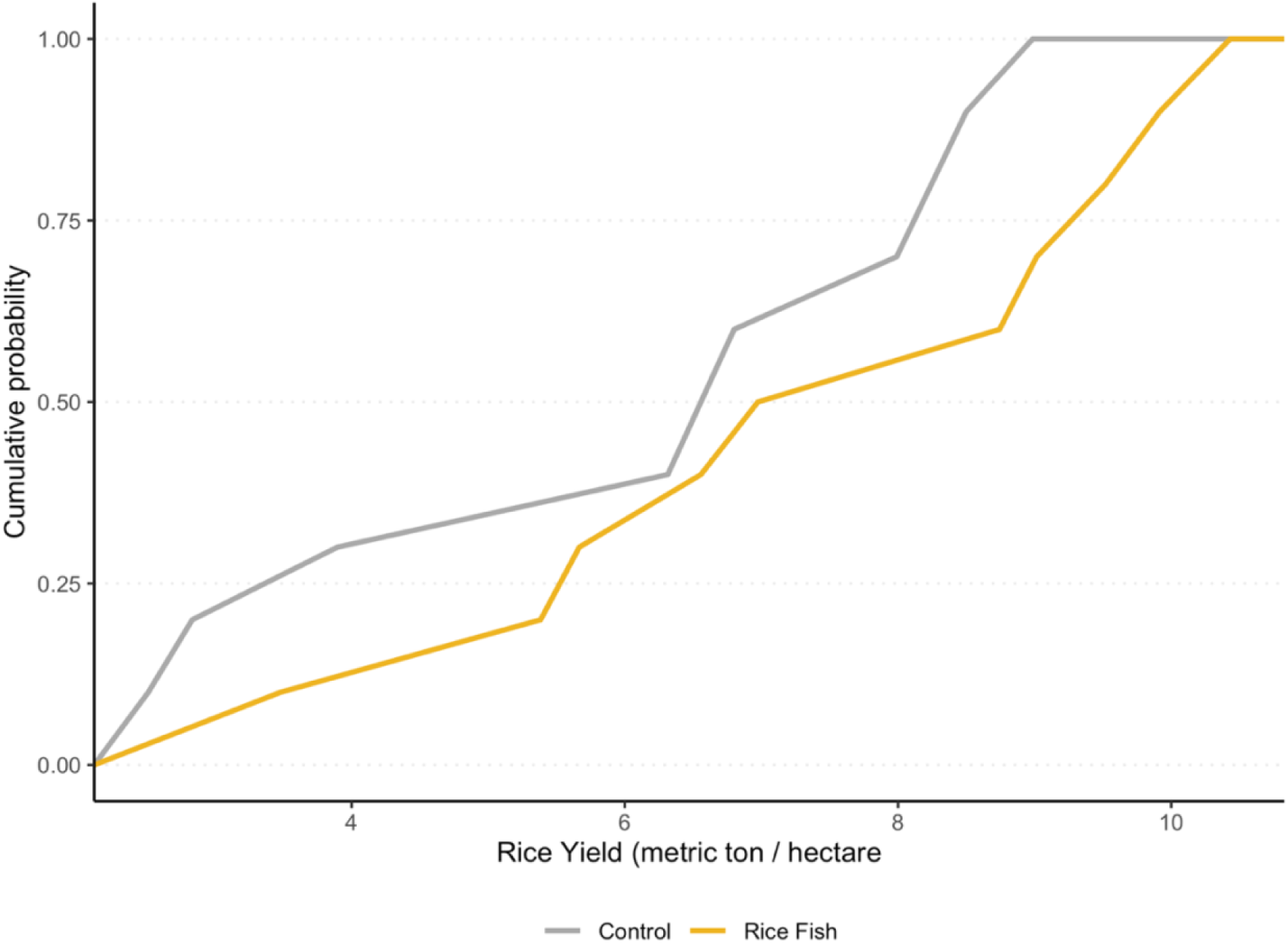
Cumulative distribution function plot of rice yield (in metric ton / hectare). The complete rightward shift in the cumulative density function for rice yields when co-cultured with fish indicates second-order stochastic dominance, meaning that any risk-averse farmer would favor rice-fish co-culture over rice monoculture in terms of rice yield. Visually, the plot suggests first-order stochastic dominance, which would mean that even risk-neutral farmers would favor rice-fish co-culture over pure stand rice cultivation, in terms of rice production alone.

Across a global meta-analysis, rice-fish systems saw a 98% increase in profitability to farmers^76^, whereas our estimates are between 334% - 949%, excluding the initial investment and additional labor requirements. Many rice-fish co-culture systems incorporate additional feed inputs for the fish^17^, which contributes nutrients to the system but increases costs. We utilized a LEISA and demonstrate nutritional improvement, high yields, and significant returns on investment. Furthermore, our estimated benefit-to-cost ratio is conservative because it does not include prospective health gains from reductions in schistosomiasis.

### Implications and Conclusions

This study demonstrates a need to target transmission of human schistosomiasis in rice fields and validates that this risk can be mitigated through a locally viable, ecologically grounded, and economically beneficial intervention: rice-fish co-culturing. By integrating native fish into rice paddies in Senegal’s semi-arid SRV, we achieved reductions in intermediate host snail abundance, increased rice and fish yields, improved soil nutrient profiles, and generated substantial economic returns for smallholder farmers. Focus group discussions were held with all farmers at each village after rice harvest to gather insight into their experiences during the trials, and most farmers expressed positive experiences and willingness to be involved in future work. These outcomes represent a rare convergence of public health protection, food security, and environmental sustainability.

As rice cultivation expands across sub-Saharan Africa—often without corresponding increases in yield or disease control infrastructure^10^—our findings carry urgent relevance. The practice of rice-fish co-culturing both addresses immediate disease transmission pathways by suppressing snail hosts and supports long-term agricultural resilience and production by reducing reliance on synthetic inputs and enhancing system productivity^10,13,18,86^. These are critical advances for a region facing intersecting pressures from climate change, sluggish agricultural productivity, poverty, malnutrition, and infectious disease.

As global demand for food and fish proteins increase, rice-fish systems offer tangible means to help keep agriculture within planetary boundaries, particularly those related to freshwater use, nutrient flows, and biodiversity loss^76,87^. A recent meta-analysis assessed the ecosystem impacts of rice-animal systems and found a reduction in nitrogen run-off into external ecosystems, reduced soil nutrient leaching, and improvements to rice yield and soil and water nutrient changes^76^. These benefits from an integrated agricultural system can have pronounced impacts at scale. Given the alignment of these outcomes with multiple Sustainable Development Goals—including no poverty, zero hunger, good health and well-being, climate action, and life below water—this intervention holds potential far beyond Senegal.

This study demonstrates that agricultural, environmental and disease control interventions need not be siloed. Our results affirm that Planetary Health approaches can deliver integrated solutions—ones that reduce disease transmission not through top-down treatment campaigns alone, but through ecologically rational redesign of food systems^2^. This is particularly salient for schistosomiasis, which is difficult to control with drug treatment programs alone^63,88^, necessitating the need for integrated interventions^47,48,89^. The incorporation of rice-fish systems could shift the paradigm from reactive treatment to proactive prevention.

We therefore call for broader implementation trials across schistosomiasis-endemic regions in sub-Saharan Africa, especially in areas where native fish species with similar ecological traits as those tested here can be employed and where rice is a major crop, as rice-associated disease risk is not isolated to Senegal^36,37,68,90,91^. Future work should also rigorously quantify the direct impacts of co-culturing on schistosome transmission and human infection outcomes, including among adult rice farmers. Our work suggests that rice-fish co-culturing is not only a tool for improved yields — it may be a scalable, systems-based strategy for breaking cycles of poverty and disease while advancing sustainable development.

## Methods

### Household and Parasitological Survey

To explore risks of schistosomiasis transmission to humans in rice agriculture settings, we leveraged an already published 2016 survey of households along the river and lake of the SRV^56,92^, focusing on eight villages, each with at least three households reporting rice agriculture (*n*=405 households). These household surveys were accompanied by individual-level parasitological data collected from school-aged children in the same year in each village (*n*=731 children). Important individual-, household-, and village-level variables were gathered in the household survey and parasitological collections. Individual-level variables included the gender and age of each child providing the parasitological sample. Household information included rice cultivation area and whether the household used piped water for laundry, as a proxy for water contact assuming that households without access to piped water have greater contact with waterbodies. Village-level covariates included total irrigated area and total fertilizer usage, both of which were calculated as the sum of households surveyed. Descriptive statistics can be found in Supplemental Table 1.

We investigated whether the presence and intensity of *Schistosoma* spp. in school-aged children living in households with rice cultivation were positively associated with the intensity of rice cultivation, controlling for several village-, household-, and individual-level variables. We used generalized linear mixed effect models (*glmmTMB* package, in *R*^93^) to explore the influence of household rice on child schistosomiasis infections with a binomial error distribution for prevalence of infection and a negative binomial error distribution for intensity of infection (count of worm egg data averaged over two testing dates for only infected children). To account for there being multiple children within households and within villages that were not completely independent, we fit random intercepts for households nested within villages. Since 3 of the 8 villages received mass drug administration of praziquantel the year prior to the survey, we used a blocking variable to control for this village-level effect in all models. Additionally, we assessed whether the correlations between rice cultivation and infection outcomes were consistent when looking across only the 5 villages which were not known to be treated the year prior to the study (see Supplemental Table 2). Reinfection rates are extremely high in the region^2^, likely explaining why the trends do not differ between analyses run on the full 8 villages versus those run on the subset of 5 villages that were not known to be treated the year prior.

### Rice-Fish Co-Culture Trials: Fish Husbandry

Tilapia were sourced from the National Agency for Agricultural Integration and Development (ANIDA) fish farm in Maraye, Senegal and maintained at Station d’Innovation Aquacole (SIA) in Saint-Louis, Senegal prior to introductions into fields. African Bonytongue below 980g were sourced from local fishermen in Mboubene, Senegal, and maintained at SIA prior to introductions. Both fish species were maintained in large, freshwater, solid-sided holding tanks (2 individuals per m^3^ tank for African Bonytongue and 1200 individuals per 30 m^3^ tank for tilapia) and fed extruded commercial fish feed. Fish were transferred to fields for stocking in a 1.5 m^3^ isotherm tank with an aeration system.

### Rice-Fish Co-Culture Trials: Experimental Design and Sampling

Two cycles of experiments of rice-fish co-culturing were performed across 4 villages of the SRV, one in 2023 (main rice cultivation season) and the other in 2024 (alternative rice cultivation season). The first trial occurred April-August 2023 in 24 paired rice fields across 2 villages. Fields were paired by farmer (when possible) such that each farmer cultivated one field of each treatment (rice monoculture or rice-fish co-culture), and the two fields were chosen to be of approximately the same size and of minimal distance to each other. When a participating farmer only had one field, two farmers’ fields were paired by size. Specifically, we tested 1) whether fish survived and grew in rice fields, 2) whether fish introductions affected rice yields, and 3) whether competitor or predator fish influenced the abundance of snails in the fields. Each of the paired fields (*n*=12), had one field designated as an unmanipulated control and the other stocked with Nile Tilapia and, when possible, African Bonytongue. Not all treatment fields could be stocked with African Bonytongue because of low numbers of fish available. 6 fields (3 pairs) were removed from rice production analyses because farmers did not use the same variety of rice in both monoculture and co-culture fields (*n*=1 pair) or reported that their fingerlings were lost to predators early on in the trial (*n*=2 pairs) (i.e., 18 paired fields were included in further analyses).

Tilapia were selected as a potential snail competitor fish because they have previously been utilized in integrated rice systems, have been successfully reared in low-density pond aquaculture^28,94^, and are known to consume algae off of rice stalks^23^. They were stocked at a density of 0.2 fingerlings/m^2^. African Bonytongue were selected because they are known snail predators in the region,^39^ have a high consumptive value and a fast growth rate, and are well suited for murky water conditions^95^, such as rice fields. African Bonytongue were stocked at a density of 2 individuals/field. Treatment fields additionally had a trench (∼1 m) dug along one side of the field to serve as a deep-water refuge for the fish and fish were stoked after the final pesticide application. Farmers were asked to apply the same level of fertilizers and pesticides to both treatment and control fields. Snail sampling was done by conducting sweep netting in the fields within 2 weeks of rice harvest. Nine 1-m sweeps were performed along the periphery of the rice field where water was accessible for sampling without damaging the crop. Three of the nine sweeps were performed in the trench in treatment fields. Depth was recorded at each sweep and all snail species were collected and counted. *Bulinus* spp. were brought back to the lab for cercarial shedding by exposing them to light for 1 hour in approximately 2 mL of DI water, utilizing standard staining and microscopy techniques^96^. At harvest, rice fields were drained. Fish were gathered into the trenches and counted and weighed by species, including any non-stocked species, except for African Bonytongue, which were individually weighed. Nile tilapia were not separated by size to calculate the average mass except for three fields, but average mass was not adjusted for fields where tilapia were not size separated (meaning that harvest size across fields underestimates growth). Rice yield information was self-reported by farmers.

The second trial occurred August-December 2024 in 13 unpaired rice fields across 3 villages (one of the same villages and farmers from the prior experiment, but not the others because the second trial was performed during the alternate cultivation season when many villages farm vegetables, instead of rice). Fields were randomized controlling for size and village (every village had at least one field of each treatment). As before, tilapia and African Bonytongue were added to rice fields after the final pesticide application in a polyculture; however, African Bonytongue were increased in density to 4 individuals/field. Tilapia were stocked in every treatment field, and African Bonytongue were stocked in 6 of the 7 treatment fields. All the methods were the same as the previous experiment with the following exceptions. Field sampling was done by sweep netting a few weeks before rice harvest at 10 points within the field along the periphery, by sweeping a 1-m area within the rice crop at 5-meter transects along 3 sides and at one corner of the field. This damaged the rice, and farmers were fully compensated for their lost crops. Insect abundance was quantified from each sweep. Nile tilapia were size-separated for weight at harvest for all fields, and only stocked-fish growth is reported.

Three 10-cm deep sub-surface soil cores were taken from one corner of each field at least half a meter back from the field edge. In treatment fields, we selected a corner that was not adjacent to the deep-water refuge. These soil cores were air dried back in the laboratory, homogenized, and then stored at room temperature in plastic bags until processing by Institut Senegalais de Recherches Agricoles (ISRA) in Saint-Louis, Senegal. ISRA used a modified Black and Wakley method to determine soil oxidized carbon, the Kjeldhal method to calculate total nitrogen, and an extraction of sodium bicarbonate and ammonium fluoride to calculate available phosphorus. Water samples were taken from three sides of the field at three points that corresponded to sweep net sampling. Treatment fields had one water sample taken from the water in the trench. Water was gathered using a syringe, filtered through a 25-mm glass microfiber filter with a 0.7 µm pore size into a 15-mL tube, frozen, and transported to the University of Notre Dame, where they were run on a Lachat Quikchem^®^ 8500 Flow Injection Analyzer by contract, as described by Mahl et al^97^, for ammonia, soluble reactive phosphorus (SRP), and nitrate.

In the second experiment, the Diama Dam was opened from September to December 2024. This reduction of water retainment in the irrigation canals caused 9 of the 13 fields to be completely dry at some point in the experiment and almost cause complete crop loss in four fields in one area of our study. Although two of those fields were able to maintain water levels in the deep-water refuge for the fish, growth was minimal given the reduced space for the fish to forage. These four fields were removed from ecological nutrient and rice production analyses. Three fields were not sampled for snails because they were dry at the time of sampling. While soil cores and water samples were taken from all fields when possible (i.e., water present in field), processing was only run on six fields for soil cores (where water was in the fields for more than two weeks prior to sampling) and for nine fields for water samples.

During the 2024 trial only, we also performed sweep net sampling for snails only in the irrigation canals adjacent to each field. Three sweeps per stretch of canal were performed, and any *Bulinus* spp. collected were taken to the laboratory for cercarial shedding. Depth was recorded at each sweep.

### Statistical Analyses of the Rice-Fish Co-Culturing Trials

All generalized linear mixed effects models were run in *R* (version 4.4.1). We used *glmmTMB* function (*glmmTMB* package, in *R*^93^) for all regression analyses. Models were compared by AIC to identify the most parsimonious model that included key covariates, and we used backwards selection to remove unnecessary covariates from our models. The quality of fit was assessed using *DHARMa* (in *R*^98^). For all significant categorical predictors from models, we calculated estimated marginal means and ran pairwise comparisons (*emmeans* package, in *R*^99^) to determine differences among categories. Figures were created using *ggplot2* and *patchwork* (in *R*^100,101^) using raw data (t tests) and using model predictions (from *ggpredict*; *ggeffects* package, in R^102^) for regression analyses.

To test whether rice yields differ between treatment and control fields, we used paired t-tests for the first trial (2023) and unpaired t-tests for the second trial (2024; two-tailed tests). To test whether tilapia and African Bonytongue weights were greater at harvest than at stocking, we used a one-tailed paired t-test. Tilapia weights were calculated as the average for each field at stocking or harvest and compared across fields, whereas for African Bonytongue each fish was individually weighed (compared across individuals in 2023 and across field averages in 2024). For significant differences, we calculated Cohen’s D effect sizes with hedge’s correction for a small sample size.

To assess differences in snail abundance in rice fields, we used generalized linear mixed models with a negative binomial error distribution. For the first trial, we tested whether the presence of African Bonytongue influenced snail abundance, because we hypothesized that African Bonytongue would consume snails in this system. Thus, we used a one-tailed pairwise test and assessed snail abundance between our two types of rice-fish co-culture fields: those stocked with African Bonytongue and tilapia and those with only tilapia. We included the number of tilapia harvested as a covariate to control for any competitive effects of large tilapia populations on snail abundance, and we controlled for village-level variation by including village as a random intercept. We assessed snail abundance at the field level for total snails and each grouping of snails (*Bulinus truncatus* and *B. globosus, B. forskalii* and *B. senegalenesis,* and *Lymnaea* spp.) separately.

For the second trial, we only had one field where tilapia, but not African Bonytongue, were stocked. Thus, we assessed snail abundance between rice-fish co-culture and control fields and did not account for the number of tilapia harvested, because control fields did not have tilapia. We included village as a random intercept and used generalized linear mixed models with a negative binomial error distribution. Because of variable water levels over the season and within a field, we tested whether water depth influenced our results by analyzing snail abundance at the sweep level, controlling for sweep depth, between co-culture and control fields while accounting for field nested within village as a random intercept.

We used generalized linear mixed effect models to assess the effect of fish additions on insect abundance and nutrients in soil and water samples, controlling for sweep depth (for insect abundance), and including field nested within village as random intercepts to account for the lack of independence of multiple samples taken within one field. For insect abundance, the error distribution was negative binomial whereas for soil and water nutrients, the distribution was Gaussian.

### Economic Analyses

We calculated the net benefit of adding fish to rice fields in northern Senegal using the first rice-fish co-culturing experiment when water levels were consistent in fields, which is typical for the region. The benefits of rice-fish co-culturing are the rice yield and the sale of harvested fish. The costs of rice-fish co-culturing are the initial cost of installing a deep-water trench and annual costs of acquiring fingerlings. We defined the cost of African Bonytongue as negligible because they were wild caught, are not available on the market, and were stocked at a density of only 2-4 individuals/field. Full details of benefits and costs are included in Supplemental Table 3.

The first-year cost of rice-fish co-culturing was estimated at $1,360–$1,540/ha, and the recurring costs thereafter at $360-$540/ha/year. Rice is valued at $0.2671/kg (Supplemental Table 3), and including fish in rice fields improved rice yields by an average of 1,793.5 kg/ha (see Figure 2). The market valuation for an individual Nile tilapia provided by the National Aquaculture Agency was 1,700 – 3,000 CFA at the time of this study. Since African Bonytongue are not readily sold at markets and they have a high non-commercial value (i.e., are often utilized for celebrations), we excluded them from net benefit calculations. However, given the extensive growth of the African Bonytongue in rice fields and their implicitly higher value because they are kept rather than sold, further efforts to include the financial benefits from the sale of African Bonytongue are recommended and likely would increase the system net-benefits. The average size of a household in the Saint-Louis region of Senegal was assumed to be 10 people based on the region’s 2018-2019 national survey^103^. In the 2016 household survey used in this study, half of the villages had households farming rice, with an average of 30.2% of households farming an average of 2.1 hectares of rice within a village. We use a FCFA to USD exchange rate of 0.0018^104^.

Using the information above and in Supplemental Table 3, and after paying for the initial cost of the deep-water trench, the net benefits/ha/year was calculated as:

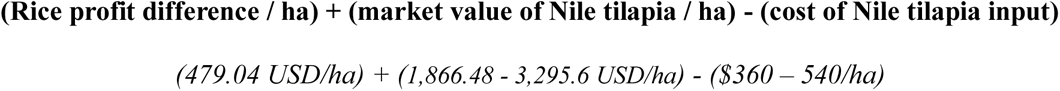

The average net benefit per rice farming household in the region was calculated as:

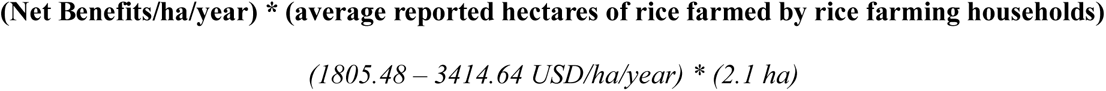

Average net benefit per capita for rice farming households of 379.16 - 717.07 USD/year.

We additionally assessed the variance of rice yields in rice monoculture versus that of rice-fish co-culture to understand how rice-fish co-culture impacts the variability of rice yields by investigating the cumulative distribution function of rice yield (metric ton/ha) and their stochastic dominance. These analyses ignore upfront investments and management for rice-fish co-culturing and consider only the variability of rice yields in these systems.

## Acknowledgements

We acknowledge the farmers and their families for their collaboration during the trials, as well as the children and families who contributed to the parasitology and survey data used in these analyses. We also thank the team at the Station d’Innovation Aquacole for their essential support and technical expertise, which were critical to the success of this work.

