## Supplemental Tables for "Rice-Fish Co-Culturing for Sustainability, Food Security, and Disease and Poverty Reduction"

**Supplemental Table 1.** Descriptive statistics of the household survey and parasitological data from children in 8 rice-farming villages in the Senegal River Basin utilized in the child occupational risk analyses.

| | **Level** | **Overall** | ***S. mansoni* positive*** | ***S. haematobium* positive** |
|---|---|---|---|---|
| **Individual-level data** | | | | |
| Children tested [*n* (%)] | | 731 | 261 (35.7) | 498 (68.1) |
| Age [*n* (%)] | *0 – 4 years* | 5 | 2 (40.0) | 3 (60.0) |
| | *5 – 6 years* | 58 | 15 (25.9) | 41 (70.7) |
| | *7 – 8 years* | 359 | 107 (29.8) | 228 (63.5) |
| | *9 – 10 years* | 218 | 90 (41.3) | 161 (73.8) |
| | *11 – 12 years* | 68 | 32 (47.1) | 47 (69.1) |
| | *13 – 16 years* | 23 | 15 (65.2) | 18 (78.3) |
| Sex [*n* (%)] | *Male* | 384 | 143 (37.2) | 269 (70.1) |
| | *Female* | 347 | 118 (34.0) | 229 (66.0) |
| Household farming participation [*n* (%)] | *Rice Farming* | 207 | 77 (37.2) | 148 (71.5) |
| | *Non-Rice Farming* | 524 | 184 (35.1) | 350 (66.8) |
| **Household-level data** | | | | |
| Households surveyed | | 405 | | |
| Households farming rice [*n* (%)] | | 115 (28.4) | | |
| Hectares of rice farmed [mean (SD)] | | 1.98 (4.7)** | | |
| Hectares of rice farmed [median (range)] | | 0.8 (0.2-39) | | |
| Households piped water usage [n (%)] | *Drinking* | 398 (98.3) | | |
| | *Laundry* | 275 (67.9) | | |
| **Village-level data** | | | | |
| Proportion households farming rice [mean (SD)] | | 30.2 (30.6) | | |
| Proportion households farming rice [median (range)] | | 12.6 (3.1-75) | | |
| Proportion households with piped water [mean (SD)] | *Drinking* | 98.6 (2.5) | | |
| | *Laundry* | 64.4 (24.9) | | |

*4 children did not have data for *S. mansoni*, so only 727 children were included in analyses for *S. mansoni* but descriptives are given out of full 731

**only 1 village has households farming more than 5 hectares of rice

**Supplemental Table 2.** Regression model outputs for the best models (selected based on lowest AIC) for the prevalence and intensity of *S. mansoni* and *S. haematobium* among children among the five villages which were not known to be provided with mass drug administration of praziquantel in 2015. Household and village were included as nested random intercepts. All log values are natural log transformed. River versus Lake was not included as a covariate because there were not enough replicates within the 5 villages.

| **Dependent variable** | **Statistical test (Error distribution)** | **Predictors** | **Chi-square** | **Estimate (SE)** | ***p* value** |
|---|---|---|---|---|---|
| Child *S. mansoni* infection | Regression (Binomial) | Child age | 6.28 | 0.2 (0.07) | 0.012 |
| | | Household hectares of rice (log) | 6.12 | 0.4 (0.1) | 0.013 |
| | | Village Irrigation Level | 4.64 | -0.02 (0.009) | 0.031 |
| | | Piped Water | 9.3 | -0.9 (0.3) | 0.0023 |
| Child *S. mansoni* infection intensity | Regression (Neg Binomial) | Household hectares of rice (log) | 0.0075 | 0.02 (0.2) | 0.93 |
| | | Village Irrigation Level | 1.19 | 0.01 (0.01) | 0.28 |
| Child *S. haematobium* infection | Regression (Binomial) | Household hectares of rice (log) | 1.44 | 0.2 (0.1) | 0.23 |
| | | Village Irrigation Level | 0.24 | 0.003 (0.007) | 0.62 |
| Child *S. haematobium* infection intensity | Regression (Neg Binomial) | Household hectares of rice (log) | 0.93 | 0.18 (0.18) | 0.33 |
| | | Village Irrigation Level | 0.79 | 0.0096 (0.01) | 0.37 |

**Supplemental Table 3.** Estimated costs and benefits utilized to calculate the annual cost-benefit ratio of rice-fish co-culturing to rice farmers.

| Cost/Benefit | Value | Estimated Value per hectare | Source | Comments |
|---|---|---|---|---|
| **Costs** | | | | |
| Nile tilapia fry | 100 – 150 FCFA /individual | 360 – 540 USD/ha | Agence Nationale de l'Aquaculture, Senegal - personnel communication | Stocking density of 2000/ha |
| Deep-water trench | | 1,000 USD/ha | SIA's records | |
| **Benefits** | | | | |
| Rice market price | 0.2671 USD/kg | | Producer Price of rice in Senegal – Food and Agriculture Organization of the United Nations | |
| Rice Profit Difference | | 479.04 USD/ha | | From an average increase of 1,793.5 kg/ha of rice when fish are added in 2023 trial data |
| Nile tilapia market price | 1700 – 3000 FCFA/individual | 1,866.48 - 3,295.6 USD/ha | Agence Nationale de l'Aquaculture, Senegal - personnel communication | Based on a mean survival of 30.8%, 616 market sized tilapia/ha, at average harvest weight of 80.59g |
